# *Streptococcus suis* encodes multiple allelic variants of a phase-variable Type III DNA methyltransferase, ModS, that control distinct phasevarions

**DOI:** 10.1101/2021.01.26.428357

**Authors:** Greg Tram, Freda E.-C. Jen, Zachary N. Phillips, Jamie Timms, Asma-Ul Husna, Michael P. Jennings, Patrick J. Blackall, John M. Atack

**Affiliations:** Institute for Glycomics, Griffith University, Gold Coast, Queensland 4222, Australia; Queensland Alliance for Agriculture and Food Innovation, The University of Queensland, St. Lucia, Queensland, 4072, Australia

**Author notes:** Address for correspondence – John M. Atack, Institute for Glycomics, Griffith University, Gold Coast, Queensland 4215, Australia.

## Abstract

*Streptococcus suis* is a significant cause of bacterial meningitis in humans, particularly in S.E. Asia, and is a leading cause of respiratory and invasive disease in pigs. Phase-variable DNA methyltransferases, associated with Restriction-Modification (R-M) systems, are a source of epigenetic gene regulation, controlling the expression of multiple genes. These systems are known as phasevarions (phase-variable regulons), and have been characterised in many host-adapted bacterial pathogens. We recently described the presence of a Type III DNA methyltransferase in *S. suis*, ModS, which contains a simple sequence repeat (SSR) tract within the open reading frame of the *modS* gene, and which varied in length between individual strains. We also observed multiple allelic variants of the *modS* gene were present in a population of *S. suis* isolates. Here, we demonstrate that a biphasic ON-OFF switching of expression occurs in the two most common ModS alleles, ModS1 and ModS2, and that switching is dependent on SSR tract length. Further, we show that ModS1 and ModS2 are active methyltransferases in *S. suis* using Single-Molecule, Real Time (SMRT) sequencing. ON-OFF switching of each ModS allele results in the regulation of distinct phasevarions, with the ModS2 phasevarion impacting growth patterns and antibiotic resistance. This is the first demonstration of a phase-variable Type III DNA methyltransferase in a Gram-positive organism that controls a phasevarion. Characterising the phenotypic effects of phasevarions in *S. suis* is key to understanding pathogenesis and the development of future vaccines.

**Importance:** *Streptococcus suis* is a causative agent of meningitis, polyarthritis and polyserositis in swine, and is a major cause of zoonotic meningitis in humans. Here we investigate epigenetic gene regulation in *S. suis* by multiple phasevarions controlled by the phase-variable Type III DNA methyltransferase ModS. This is the first characterised example of a Type III R-M system regulating a phasevarion in a Gram-positive organism. We demonstrate that biphasic ON-OFF switching of ModS expression results in differences in bacterial growth and antibiotic resistance. Understanding the effects of ModS phase variation is required to determine the stably expressed antigenic repertoire of *S. suis*, which will direct and inform the development of antimicrobial treatments and vaccines against this important pathogen.

## Introduction

*Streptococcus suis* is an important veterinary pathogen that contributes a substantial disease burden to the swine industry (1). *S. suis* is also a significant pathogen in humans, causing zoonotic meningitis, most commonly associated with occupational exposure (2) and consumption of contaminated pork products (3). *S. suis* frequently colonises the upper respiratory tract of pigs, where it is considered a commensal organism (4). Virulent strains of *S. suis* are able to adhere to and invade epithelial cells of the airway allowing access to the bloodstream. In both pigs and humans, *S. suis* is able to breach the blood brain barrier, resulting in the development of meningitis and septic shock (4).

Phase variation is the rapid and reversible switching of gene expression, and plays an important role in the pathogenesis of many organisms, where it is usually associated with expression of surface factors such as capsules (5), adhesins (6, 7) and lipooligosaccharide biosynthesis (8). Randomised gene expression as a result of phase variation can complicate vaccine development as it results in an unstable antigenic repertoire, and is potentially a major cause of vaccine escape. Many bacterial pathogens also encode cytoplasmically located phase-variable DNA methyltransferases, that are associated with restriction-modification (R-M) systems (9). Variable methyltransferase expression results in genome wide methylation differences, resulting in differential gene regulation by epigenetic mechanisms in systems called phasevarions (phase variable regulons) (9, 10). Many phasevarions are controlled by the ON-OFF switching of Type III DNA methyltransferases, encoded by *mod* genes. Phasevarions controlled by Type III *mod* genes have been described in *Haemophilus influenzae* (11, 12), *Moraxella catarrhalis* (13), *Neisseria* spp. (14), *Helicobacter pylori* (15) and *Kingella kingae* (16). The phasevarions in these human adapted pathogens all control expression of genes involved in the pathogenesis of these organisms (17–19). All these phase variable *mod* genes contain simple DNA sequence repeat (SSR) tracts within their open reading frame. These SSRs tracts are highly unstable and prone to variation in length due to slipped-strand mispairing during DNA replication, resulting in the biphasic ON-OFF switching of gene expression; the *mod* gene is either in-frame and Mod is expressed (ON), or a variation in SSR tract length results in a frameshift and premature stop codon, and Mod is not expressed (OFF) (20).

The methyltransferase specificity of a Type III Mod protein is dictated by the central target recognition domain (TRD) of the encoding *mod* gene (9, 10). This TRD varies between the alleles of individual *mod* genes, with the 5′ and 3′ regions of individual *mod* genes being highly conserved between alleles (9, 10). For example, there are twenty-one *modA* alleles encoded by non-typeable *Haemophilus influenzae* (NTHi) and *Neisseria* species; six *modB* and seven *modD* alleles have been identified in *Neisseria gonorrhoeae* and *Neisseria meningitidis*; and nineteen *modH* alleles are present in *H. pylori* (10, 21). Our previous analysis of the restriction enzyme database REBASE demonstrated the presence of a *mod* gene in *S. suis*, which we have named *modS*, that contained a GAGCA_(n)_ SSR tract (22). A follow up study analysing of a large collection of *S. suis* isolates determined the presence of three distinct *modS* alleles, each of which methylated an adenine in distinct DNA sequences. ModS1 methylated 5′-GCG^(m6)^**A** DT-3′ (D is either A,G or T), ModS2 methylated 5′-VTC^(m6)^**A** TC-3′ (V is either A,G or C), and ModS3 (present in a single strain identified in GenBank) methylated 5′-GTTC^(m6)^**A** NNNB-3′ (B is either C, G or T; N is any nucleotide) (23). These specificities were determined through heterologous expression of the methyltransferase in *E. coli*. Whilst variable length GAGCA_(n)_ SSR tracts were present in different strains of *S. suis* encoding the same *modS* allele, it was not determined whether *modS* is phase variable, or whether the ModS protein is an active methyltransferase in *S. suis* when expressed.

In this study we investigated phase variation of the two most prevalent *modS* alleles, *modS1* and *modS2*, by enriching populations of *S. suis* strains for specific GAGCA_(n)_ SSR tract lengths in each encoding *modS* gene. We then determined if differential protein expression occurred in enriched ON-OFF populations of *S. suis* using SWATH-MS. Clinically relevant phenotypes were assessed *in vitro* to determine if phase variation of the ModS methyltransferase could result in relevant phenotypic changes or advantages *in vivo*. Studying the impact of these systems provides an understanding of not only disease pathogenesis, but will also guide the development of vaccines by defining the stably expressed antigenic repertoire of *S. suis*.

## Results

### ModS is a phase variable DNA methyltransferase

In all previous examples of phase-variable Type III *mod* genes, SSR tract length variation results in a biphasic ON-OFF switching of expression. In order to determine if variation in length of the GAGCA_(n)_ repeat tract (Figure 1A) located in the *modS* open reading frame led to phase variable expression of the gene, isogenic strains were generated containing defined lengths of this SSR tract. The isolation of three consecutive repeat tract lengths would theoretically result in one strain where the respective *modS* gene is ON, (GAGCA_(n)_ repeat tract length will place the gene in frame and expressed), and two strains of the triplet will be OFF (GAGCA_(n)_ repeat tract length will place the gene out of frame and not expressed). These enriched populations were generated in *S. suis* strains LSS89 (*modS1*) or SS1056 (*modS2*), as our analysis of the TRDs of these two alleles showed they were highly variable, and therefore predicted to methylate different target sequences (Figure 1B) as we demonstrated with heterologous over-expression of ModS1 and ModS2 previously (23). Strains were enriched for SSR tract lengths containing either 19, 20 or 21 repeats in *modS1* and 17, 18 or 19 repeats in *modS2.* Fragment analysis of each strain confirms enrichment of each SSR tract length to above 80% (Figure 1C).

**Figure 1.**
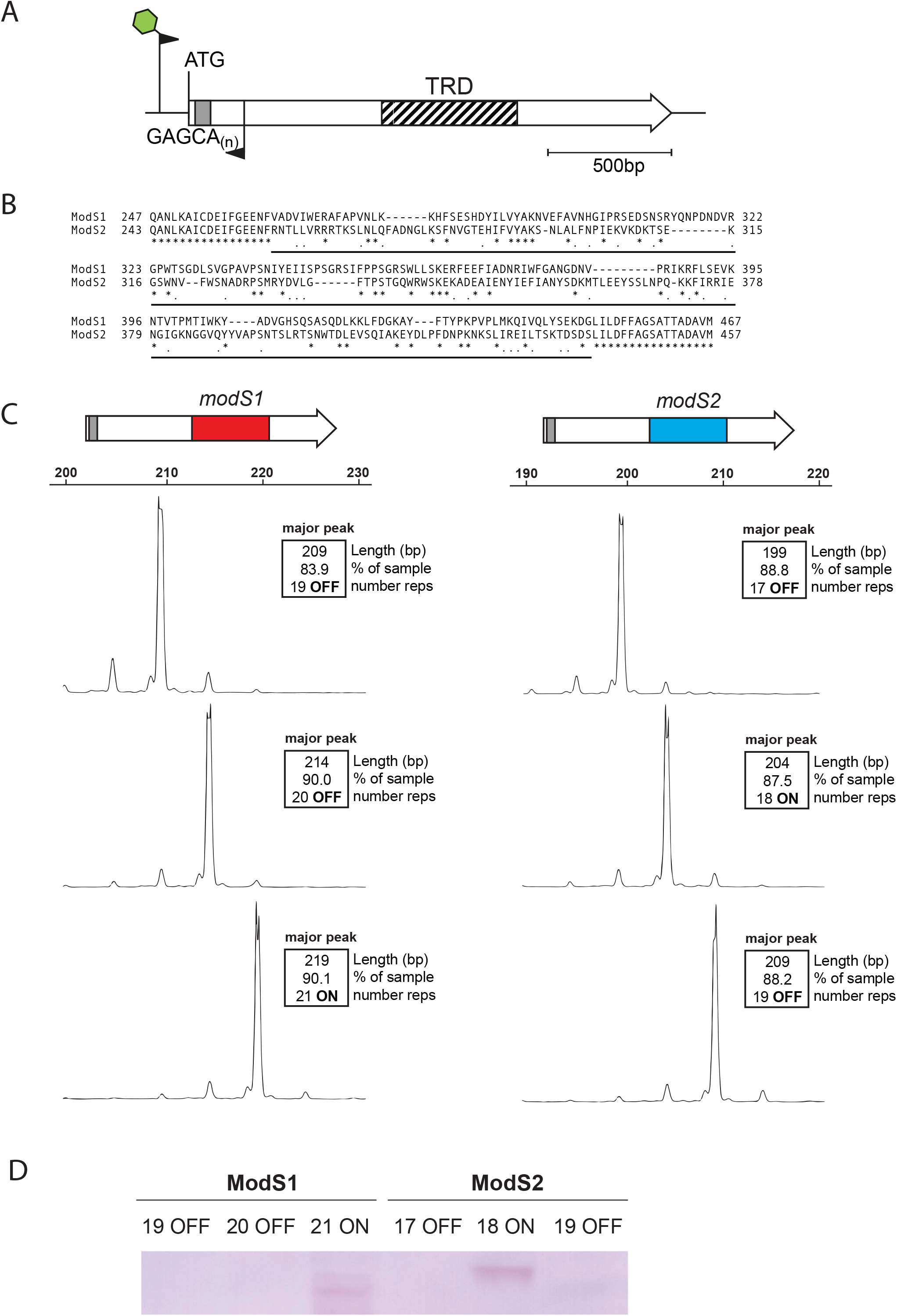
Expression of ModS alleles. **A)** The *modS* gene contains a variable length GAGCA_(n)_ SSR tract (grey box) near the start of the gene, and a variable central target recognition domain (TRD) represented by the hatched box. The 5′ and 3′ regions of the modS gene are highly (>95% nucleotide identity) conserved (white). PCR over the SSR tract was determined by FAM labelled PCR using primers SsuT3-F-FAM and SsuT3-R, and analysed using fragment length analysis; **B)** Alignment of the TRD regions of ModS1 and ModS2 showing <25% amino acid identity. * represents identical amino acid residues, ̇ represents similar amino acid residues (basic, acidic, neutral), TRD region underlined. Alignments carried out in ClustalW; **C)** Fragment length analysis traces of the enriched *modS1* and *modS2* populations of strains LSS89 and SS1056, respectively, containing three consecutive GAGCA_(n)_ SSR tract lengths; **D)** Western blot analysis using ModS antisera demonstrates that the ModS protein is only present in *S. suis* populations enriched for 21 repeats in ModS1 (LSS89) and 18 repeats in ModS2 (SS1056), demonstrating phase-variable expression of this protein.

Western blotting of whole cell lysates of these enriched strains with antisera generated against the conserved region of all ModS proteins demonstrate that variation in SSR tract length led to a biphasic ON-OFF switching of expression (Figure 1D; full blot in Supplementary Figure 1). The ModS1 protein was only produced in the LSS89 strain which contained 21 GAGCA repeats in the *modS1* gene. ModS2 was only produced in the SS1056 strain enriched for 18 GAGCA repeats. This matches the prediction from the annotated start codon of both genes (Supplementary Figure 2).

To demonstrate methyltransferase activity of the expressed ModS protein, Pacific Biosciences (PacBio) Single-Molecule, Real-Time (SMRT) sequencing was carried out on genomic DNA isolated from our triplet sets of enriched isogenic strains (19, 20, 21 repeats for *modS1* in strain LSS89; 17, 18, 19 repeats for *modS2* in strain SS1056). We detected a separate, distinct Type I methyltransferase motif in strain LSS89 compared to strain SS1056 (Table 1), with these motifs being fully methylated in all three strains of each enriched triplet set. When comparing the methylomes of enriched LSS89 strains, we detected a Type III methyltransferase motif, 5′-GCG^(m6)^**A**T, only present in the strain where ModS1 was expressed (21 repeats ON; Table 1). This methylation by ModS1 occurred at 2859 5′-GCG^(m6)^**A**T sites throughout the genome, and was not detected in strains enriched for 19 or 20 GCACA repeats in the *modS1* gene. SS1056 enriched strains exhibited a different Type III motif, 5′-VTC^(m6)^**A**TC, which was only present when ModS2 was expressed (18 repeats ON; Table 1). This motif was methylated 4810 times in the SS1056 genome, and again was not methylated in strains enriched for 17 or 19 repeats (OFF) in the *modS2* gene. These motifs are the same as detected previously when these ModS alleles were heterologously over-expressed in *E. coli* (5).

**Table 1.** Summary of methylomes for *S. suis* strains LSS89 (ModS1) and SS1056 (ModS2). Strains were enriched for GAGCA_(n)_ SSR tracts of 19, 20, or 21 GAGCA repeats in the *modS1* gene in strain LSS89, or 17, 18, or 19 GAGCA repeats in the *modS2* gene of strain SS1056. Type III methyltransferase motifs are only detected in strains enriched for an ON number of repeats (ModS1 in red in strain LSS89 21 repeats; ModS2 in blue in strain SS1056 18 repeats) and matches the protein expression detected in Figure 1. In ModS2, V represents A, G or C. % values represent motifs detected/motifs present. Full methylome data is presented in Supplementary Data 1.

|  | **LSS89 *modS1*** |  |  | **SS1056 *modS2*** |  |  |
| --- | --- | --- | --- | --- | --- | --- |
| motif | 19 reps | 20 reps | 21 reps | 17 reps | 18 reps | 19 reps |
| 5'-G<sup>(m6)</sup>AAGNNNNNTTC/<br>5'-G<sup>(m6)</sup>AAHNNNNCTTC | 100%/<br>100% | 100%/<br>100% | 100%/<br>100% |  |  |  |
| 5'-GCG<sup>(m6)</sup>AT (ModS1) | ND | ND | 99.9% |  |  |  |
| 5'-GYT<sup>(m6)</sup>ANNNNNNNNTTC/<br>5'-GA<sup>(m6)</sup>ANNNNNNNNTARC |  |  |  | 100%/<br>100% | 100%/<br>99.6% | 100%/<br>100% |
| 5'-VTC<sup>(m6)</sup>ATC (ModS2) |  |  |  | ND | 99.9% | ND |

### Biphasic switching of ModS1 and ModS2 results in the expression of distinct phasevarions

Quantitative proteomics was carried out to determine if distinct protein expression profiles occurred as a result of ModS expression in *S. suis*, i.e., if these phase-variable methyltransferases control phasevarions. The expression profiles of an ON-OFF ModS1 pair from strain LSS89 (21 repeats ON and 19 repeats OFF) and an ON-OFF ModS2 pair from strain SS1056 (18 repeats ON and 17 repeats OFF) were assessed using SWATH-MS. Both strain pairs exhibited unique changes in protein expression dependent on phase variation of their respective *modS* gene, with SWATH-MS covering ~23% of total proteins in LSS89 strains and ~22% of total in SS1056 strains. Significant differences in protein abundance between the ON and OFF populations are represented as volcano plots in Figure 2. In strain LSS89, ModS1 ON resulted in higher expression of several proteins involved in a range of cellular processes compared to strains where ModS1 was OFF. Multiple transcriptional regulators were upregulated by expression of ModS1 such as the replication initiator protein DnaA (24), and a YlbF/ YmcA type protein (25), as well as transcription factors and repressors. Many ribosomal subunits were upregulated as well as those involved in general metabolism such as a nitroreductase (26), glucokinase (27) and a protein involve in alkylphosphonate utilisation (28) (Table 2). Conversely, in strains where ModS1 was not expressed, proteins such as ribosomal proteins, several components of ABC transporters, and the fatty acid binding protein DegV (29) were all increased in expression (Table 2).

**Figure 2.**
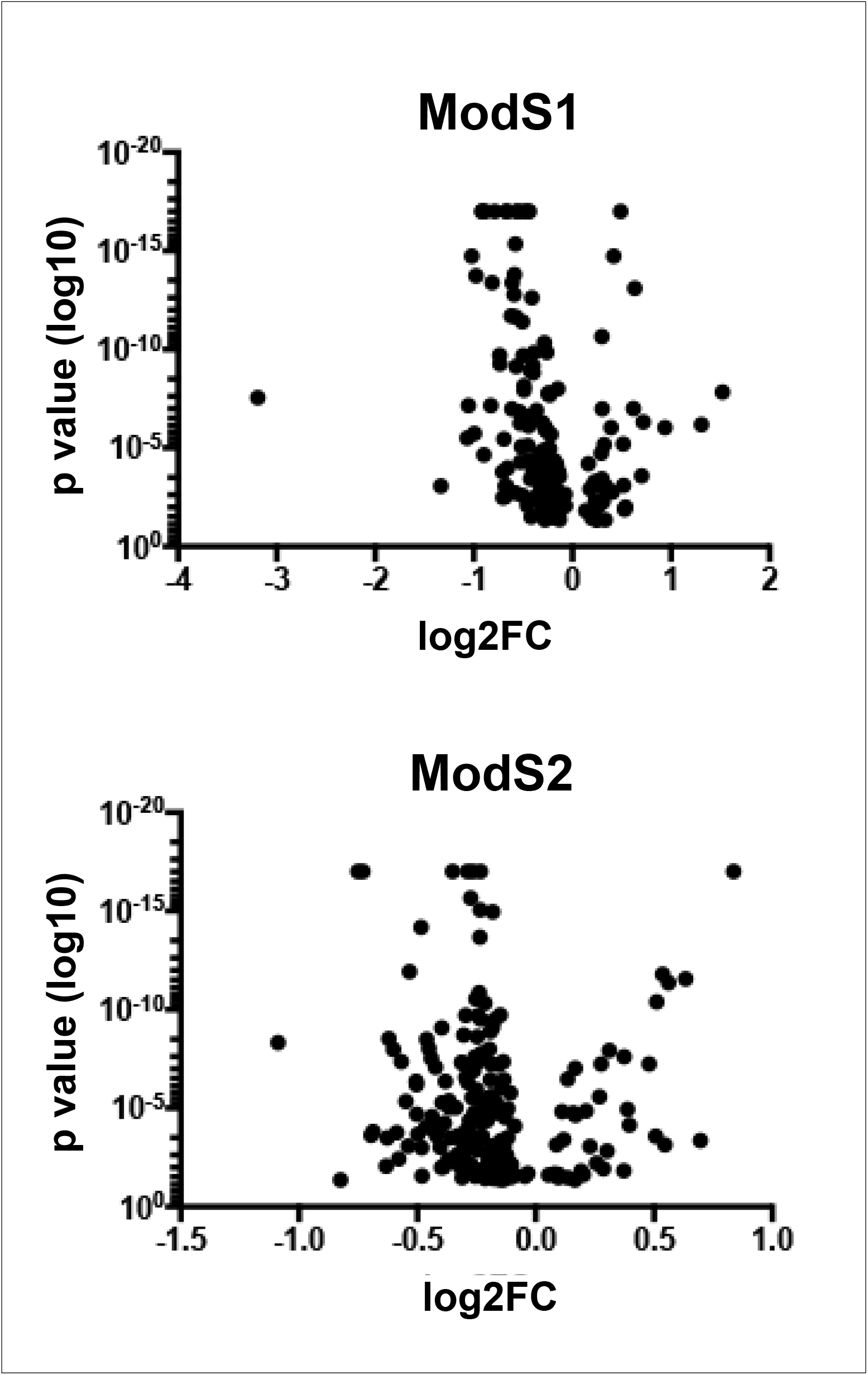
Volcano plot demonstrating changes to protein expression as a result of ModS. SWATH-MS proteomics demonstrated a coverage of 450 of 1964 identified proteins (~23%) in LSS89 ModS1 ON-OFF strain pair, and 411 of 1905 (~22%) identified proteins in SS1056 ModS2 ON-OFF strain pair. The x axis indicates relative fold difference in protein abundance in ON compared to OFF; the y axis indicates statistical significance.

**Table 2.** Differentially regulated proteins (>1.5 fold) in the ModS1 phasevarion. Fold change presented as *modS1* ON vs *modS1* OFF

| Down regulated in *modS1* ON |  |  |  |
| --- | --- | --- | --- |
| Accession | Protein | Fold change<br>(ON vs OFF) | *p*-value |
| WP_002940030.1 | 30S ribosomal protein S12 | 2.87 | 1.43E-8 |
| WP_002938143.1 | ABC transporter permease | 2.48 | 6.54E-7 |
| WP_009909754.1 | 50S ribosomal protein L27 | 1.92 | 9.04E-7 |
| WP_014638653.1 | ABC transporter substrate-binding protein | 1.65 | 4.81E-7 |
| WP_002936253.1 | DegV family protein | 1.62 | 2.53E-4 |
| WP_023369127.1 | Sugar ABC transporter substrate-binding protein | 1.55 | 7.86E-14 |
| WP_004195491.1 | 50S ribosomal protein L23 | 1.54 | 9.68E-8 |
| Up regulated in *modS1* ON |  |  |  |
| WP_002936659.1 | 50S ribosomal protein L29 | 9.14 | 2.76E-8 |
| WP_002940438.1 | Amino acid ABC transporter ATP-binding protein | 2.1 | 8.50E-4 |
| WP_012027640.1 | Ribosome biogenesis GTPase Der | 2.08 | 3.12E-6 |
| WP_002942806.1 | CsbD family protein | 2.03 | 6.94E-8 |
| WP_014636557.1 | Transcriptional repressor | 1.99 | 1.78E-15 |
| WP_002936247.1 | HU family DNA-binding protein | 1.97 | 1.81E-6 |
| WP_002936483.1 | 50S ribosomal protein L7/L12 | 1.89 | 1.82E-14 |
| WP_002938891.1 | Acyl carrier protein | 1.88 | ≤1.00E-17 |
| WP_002936048.1 | DUF1846 domain-containing protein | 1.86 | ≤1.00E-17 |
| WP_014736429.1 | Nitroreductase family protein | 1.77 | 2.17E-5 |
| WP_014735259.1 | 30S ribosomal protein S10 | 1.76 | ≤1.00E-17 |
| WP_002940682.1 | 30S ribosomal protein S16 | 1.73 | 6.91E-8 |
| WP_002934959.1 | DUF1149 family protein | 1.67 | 4.09E-14 |
| WP_023368958.1 | Chromosomal replication initiator protein DnaA | 1.66 | ≤1.00E-17 |
| WP_023369777.1 | dTDP-4-dehydrorhamnose reductase | 1.64 | 1.96E-10 |
| WP_002936660.1 | 50S ribosomal protein L16 | 1.62 | 5.40E-10 |
| **WP_004194840.1** | DUF2829 domain-containing protein | 1.6 | 1.60E-4 |
| **WP_012027178.1** | PTS sugar transporter subunit IIA | 1.6 | 3.27E-3 |
| **WP_002936656.1** | 50S ribosomal protein L24 | 1.59 | 3.52E-6 |
| **WP_002937025.1** | Response regulator transcription factor | 1.59 | 2.89E-3 |
| **WP_011922043.1** | DivIVA domain-containing protein | 1.59 | 8.40E-4 |
| **WP_002938966.1** | YlbF/YmcA family competence regulator | 1.58 | ≤1.00E-17 |
| **WP_012775074.1** | Alkylphosphonate utilization protein | 1.54 | 8.72E-4 |
| **WP_023370402.1** | ROK family glucokinase | 1.53 | ≤1.00E-17 |
| **WP_002936622.1** | 30S ribosomal protein S13 | 1.51 | 1.00E-4 |
| **WP_023369315.1** | Glycine cleavage system protein H | 1.5 | 1.94E-12 |

In strain SS1056, ModS2 ON-OFF switching resulted in varied expression of a distinct set of proteins (Table 3). ModS2 expression increased the levels of proteins involved in amino acid metabolism such as cysteine synthase and aminopeptidase D (30). Proteins involved in general metabolism also showed increased expression in ModS2 ON, such as dihydroxyacetone kinase (31). The DNA biosynthesis enzymes ribonucleotide-diphosphate reductase and ribonucleotide-triphosphate reductase (32) were increased in expression in ModS2 ON, as was an acyl carrier protein involved in fatty acid biosynthesis (33). Three proteins were upregulated in strains which did not express ModS2 (OFF); an ATP-dependent Clp protease, the nucleotide exchange factor GrpE and a glyoxalase/bleomycin resistance/extradiol dioxygenase family protein involved in resistance to antimicrobials.

**Table 3.** Differentially regulated proteins (>1.5 fold) in the ModS2 phasevarion. Fold change presented as *modS2* ON vs *modS2* OFF

| Down regulated in *modS2* ON |  |  |  |
| --- | --- | --- | --- |
| Accession | Protein | Fold change<br>(ON vs OFF) | *p*-value |
| WP_002935876.1 | Glyoxalase/bleomycin<br>resistance/extradiol dioxygenase<br>family protein | 1.79 | ≤1.00E-17 |
| WP_044673510.1 | ATP-dependent Clp protease ATP-binding subunit | 1.62 | 4.35E-4 |
| WP_024387015.1 | Nucleotide exchange factor GrpE | 1.55 | 2.81E-12 |
| Up regulated in *modS2* ON |  |  |  |
| WP_012028296.1 | 50S ribosomal protein L20 | 2.13 | 4.82E-9 |
| WP_012027287.1 | Peptidylprolyl isomerase | 1.77 | 4.40E-2 |
| WP_002938891.1 | Acyl carrier protein | 1.68 | ≤1.00E-17 |
| WP_044754372.1 | Cysteine synthase A | 1.66 | ≤1.00E-17 |
| WP_002936486.1 | 50S ribosomal protein L10 | 1.62 | 2.35E-4 |
| WP_044680965.1 | Class 1b ribonucleoside-<br>diphosphate reductase subunit<br>alpha | 1.61 | 1.54E-4 |
| WP_044771061.1 | Dihydroxyacetone kinase subunit L | 1.55 | 8.68E-3 |
| WP_024384331.1 | Aminopeptidase P family protein | 1.54 | 3.24E-4 |
| WP_044681922.1 | Ribonucleoside-triphosphate<br>reductase | 1.54 | 2.97E-9 |
| WP_079394259.1 | Amino acid ABC transporter<br>substrate-binding protein | 1.52 | 1.07E-8 |
| WP_044674536.1 | Winged helix-turn-helix<br>transcriptional regulator | 1.5 | 1.70E-4 |

### ModS switching results in differences in growth

Our proteomic analysis of ModS switching demonstrated several proteins which could potentially impact growth rate exhibited differential regulation in both ModS alleles. These included proteins involved in general metabolic enzymes, DNA transcription factors and repressors as well as biosynthesis enzymes (Tables 2 & 3). In order to determine if these altered protein expression levels could affect *S. suis* growth rates, we conducted standard growth curves of our enriched ModS ON-OFF pairs in rich media. These growth curves showed a small but repeatable (n = 3) difference in growth rate for both strain pairs (Figure 3): when ModS1 was ON, growth was to a lower final OD (P value <0.05), and conversely, when ModS2 was ON, growth was to a higher final OD (P Value <0.05).

**Figure 3.**
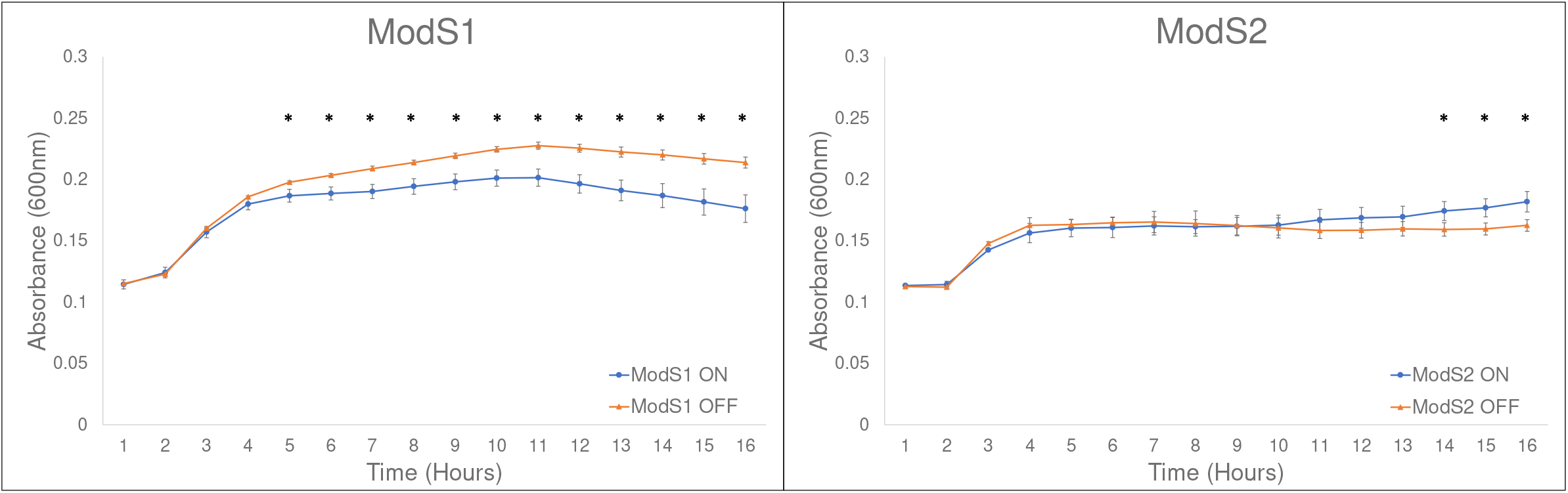
Growth Curves of *S. suis* populations enriched for *modS1* and *modS2*. ON-OFF strain pairs for ModS1 (strain LSS89) and ModS2 (SS1056) were grown in rich media (THB-Y broth) for 18 hours with shaking. Statistically significant differences (*P* value <0.05) in absorbance at each time point are indicated by asterisks, assessed using Student’s t-test.

### ModS2 phase variation results in differences in antibiotic resistance

SWATH-MS also demonstrated differential expression of an antibiotic resistance protein in ModS2 (WP_002935876.1), annotated as a glyoxalase/bleomycin resistance/extradiol dioxygenase family protein, which exhibited higher expression in strains which did not express ModS2 (OFF) compared to ModS2 ON. We therefore carried out a minimum inhibitory concentration (MIC) analysis using several beta lactam antibiotics (ampicillin, penicillin and amoxicillin) and a glycopeptide antibiotic (vancomycin) (Table 4). Although there was no change in MIC to amoxicillin, vancomycin and penicillin, there was a two-fold increase in resistance to ampicillin when ModS2 was OFF compared to ModS2 ON (0.32 μg/mL vs 0.16 μg/mL respectively).

**Table 4.** MICs of ModS2 ON vs ModS2 OFF enriched *S. suis* strains.

|  | ModS2 ON | ModS2 OFF |
| --- | --- | --- |
| **Ampicillin (µg/mL)** | **0.16** | **0.31** |
| **Vancomycin (µg/mL)** | 1.25 | 1.25 |
| **Amoxicillin (µg/mL)** | 0.16 | 0.16 |
| **Penicillin (µg/mL)** | 0.63 | 0.63 |

## Discussion

Over the last 15 years, phasevarions (phase-variable regulons) controlled by ON-OFF switching of Type III *mod* genes have emerged as an important gene regulation strategy in a diverse range of host-adapted pathogens (9, 12, 13, 15, 16, 21). In all these examples, phase variable ON-OFF switching of *mod* expression occurs via changes in the length of locus encoded SSR tracts. In this study, using a combination of strain enrichment, Western blotting, and SMRT sequencing, we conclusively demonstrate that *S. suis* contains multiple allelic variants of a new phase-variable Type III *mod* gene, *modS*, with this representing the first example of a phasevarion controlled by biphasic ON-OFF switching of a Type III DNA methyltransferase in a Gram-positive organism (Figure 1). The antisera used to detect ModS was generated against the conserved regions of ModS, and should recognise all ModS alleles with equal affinity, as sequence variation of Mod proteins only occurs in the central TRD region (9, 34). Both ModS alleles were only expressed in strains with specific SSR tract lengths; ModS1 was only present in strains with 21 GAGCA repeats and ModS2 was only expressed in strains with 18 GAGCA repeats. In both cases, the repeat length placed the *modS* gene in the correct reading frame and demonstrates that the expression of the ModS protein is dependent on the SSR tract length.

SMRT sequencing was used to determine methyltransferase activity and specificity of ModS1 and ModS2. LSS89 strains of *S. suis* enriched for ModS1 expression (21 GACAC repeats ON) exhibited methylation of the adenine in 5′-GCG^m6^**A**T motifs which was not seen in strains in which ModS1 was OFF (19 or 20 GACAC repeats). ModS2 expressing strains (SS1056) of *S. suis*, exhibited methylation of a Type III Mod motif of 5′-VTC^m6^**A**TC motifs (where V can be either A, G or C) when ModS2 was predicted to be ON (18 GACAC repeats). This confirmed our previous findings when we over-expressed ModS1 and ModS2 using a recombinant *E. coli* system (23). SMRT sequencing also demonstrates that both ModS1 and ModS2 are active methyltransferases in *S. suis*, methylating at thousands of sites in their respective genomes (in strain LSS89, 2861 5′-GCG^m6^**A**T sites are present; in SS1056, 4812 5′-TC^m6^**A**TC sites are present; Supplementary Data 1).

SWATH-MS proteomic analysis confirmed that the altered methylation resulting from ModS phase-variation results in a change in the expression of numerous proteins in distinct phasevarions. These changes in protein expression were seen across many families of protein, including those involved in central metabolism, gene regulation, and transporters. Phase-variable switching of ModS resulted in differences in growth rate, but it is unclear whether this is a result of an alteration of regulatory genes due to ModS switching, or another, as yet uncharacterised effect of ModS phase variation. Although the final differences in final OD between enriched ON and OFF strain pairs were small, they are statistically significant (P value <0.05), and could have an effect on both long-term carriage and disease, particularly if one population has an advantage resulting from differential protein expression over time. ModS2 phase variation also resulted in a difference in antibiotic resistance. A protein repressed by ModS2 expression, annotated as a glyoxalase, bleomycin resistance and extradiol dioxygenase family protein, described as involved in resistance to beta-lactam and glycopeptide antibiotics (35), which are commonly used to treat bacterial infections in humans as well as commercial pig farms (36). The MIC of *S. suis* was assessed against three commonly used beta lactams (ampicillin, amoxicillin and penicillin) and a glycopeptide from the same antibiotic class as bleomycin (vancomycin). A strain in which ModS2 was OFF, where the expression of this resistance protein was increased, showed a small (2-fold) increase in MIC towards ampicillin, compared to the isogenic strain in which ModS2 was ON. Although small, there may be a cumulative effect, particularly as many beta lactam antibiotics are extensively used prophylactically in commercial pig farms (37–40). This has the potential to result in a long-term increase in ampicillin resistance in *S. suis* populations, although this would need to be studied in a model system and is beyond the scope of this work.

Although many of the changes in protein expression determined by our SWATH-MS analysis of enriched ON-OFF population in each ModS allele were small (1.5-fold to 2-fold), SWATH-MS is a technique which quantitates protein expression based on abundance (41). Differential expression of proteins will therefore only identify differences in highly expressed proteins, and expression differences of proteins beyond the limit of detection will be missed. We recognise this is a limitation of our approach, but a SWATH-MS proteomics approach has been previously used to characterise phasevarion mediated changes to protein expression (13, 42), with similar small differences in expression also reported in these phasevarions. Therefore, whilst we have determined that both ModS1 and ModS2 affect expression of small distinct sets of proteins, it is likely that further changes to gene and protein expression remain undetected. In order to thoroughly characterise each phasevarion, further analysis will be needed. It will also be important to characterisation the phasevarions controlled by other *modS* alleles; in previous work we showed that a third *modS* allele – *modS3* – was present in *S. suis*. The ModS3 allele methylated a distinct sequence to ModS1 and ModS2 (23), and is therefore highly likely to control a different phasevarion. The presence of additional *modS* alleles should also be determined, and studied, in order to detail all gene and protein expression changes mediated by phasevarions in *S. suis*.

This characterisation of ModS is the first described instance of a phasevarion in a Gram-positive organism controlled by a Type III methyltransferase. Different alleles of ModS methylate distinct motifs and result in distinct phasevarions. These distinct ModS phasevarions result in gross differences in growth rate, and in the case of ModS2, differential resistance to antibiotics. Both these phenotypes could affect disease and pathology, and this remains to be studied using both *in vitro* models and ideally, an *in vivo* challenge using the natural host (pigs). A thorough understanding of phase variation of gene expression, and in particular phasevarions, is required in order to determine the stably expressed antigenic repertoire of a bacterial species. The prevalence of phase variable methyltransferases across the bacterial domain demonstrates that phasevarions are a widespread contingency strategy (22, 43, 44), and that characterisation of these systems is imperative in order to rationally design effective vaccines that only target stably expressed antigens in the organisms where they are present.

## Materials and methods

### Bacterial strains and growth

*S. suis* strains LSS89 and SS1056 (45) used in this study were grown in THB broth (Oxoid) supplemented with 2% yeast extract (THB-Y) or on THB-Y plates (THB-Y broth supplemented with 1.5% w/v agar). Cultures were incubated overnight at 37°C.

### Strain enrichment

Fragment length analysis of the SSR tract in each *modS* allele was conducted to determine the length of the GAGCA_(n)_ SSR tract. A PCR was performed using a fluorescently labelled forward primer SsuT3-F-FAM (5′-FAM-CAT CAA AAA CGG CTT GAC AGC C) and the reverse primer SsuT3-R (5′-GCA ATG TTG TCT GAT AAA ACA TCT TTT G) as described previously (23). DNA fragment length analysis was carried out at the Australian Genome Research Facility (AGRF; Brisbane, Australia). This technique was used to enrich populations of *S. suis* strains for defined SSR lengths through a combination of fragment length analysis and subculturing. *S. suis* populations of strain LSS89 (encoding *modS1*) were enriched for 19, 20 and 21 GAGCA repeats, and strain SS1056 (encoding modS2) were enriched for 17, 18, 19 GAGCA repeats. Populations containing >80% of each single tract length were considered to be enriched, and were used in subsequent studies.

### SDS-PAGE and Western blot

Cell lysates of each enriched strain were prepared from overnight liquid cultures of *S. suis* grown in THB-Y broth. Samples were normalised to O.D._600_ 30.0, pelleted at 12,000xg for 1 minute and resuspended in RIPA buffer (150 mM NaCl, 1% Triton X-100, 0.5% SDS, 50 mM Tris HCl, pH 8). Samples were sonicated twice for 30 seconds at a probe intensity of 15 amplitude microns, on ice. Lysates were clarified by centrifugation at 14,000xg for 20 minutes to remove cellular debris.

Supernatants were prepared for SDS-PAGE using BOLT 4x loading dye containing 100 mM dithiothreitol (Sigma) and boiled for 30 minutes, then loaded on precast 4-12% BOLT Bis-Tris gel (Thermo Fisher) and run for 45 minutes at 165V in MOPs 1x running buffer according to manufacturer’s instructions (Thermo Fisher). Proteins were visualised by Coomassie brilliant blue staining. Twice the amount of lysate was loaded onto gels for use in Western blotting and proteins were transferred to pre-equilibrated polyvinylidene difluoride (PVDF) membranes (BioRad) at 20 volts for 60 minutes using the BOLT transfer system (Thermo Fisher). Following transfer, membranes were blocked with 10% skim milk in TBS-T for 60 minutes. Membranes were incubated with a 1:1,000 dilution of anti-ModS antisera (raised as detailed below) overnight at 4°C, followed by a 1:10,000 anti-mouse alkaline-phosphatase secondary antibody (Sigma). Blots were developed using NBT/BCIP (Roche) according to manufacturer’s instructions.

### Construction of a TRD-less ModS protein

The *modS1* gene was amplified from *S. suis* strain LSS89 excluding the CAGCA_(n)_ SSR tract using primers SsuT3-oE-F (5′-AGTCAG <u>CATATG</u> AGC AGA GCA AAG CAA AAG CTT GGA GAA TAC ACT CAA G) and SsuT3-oE-R (5′-AGTCAG <u>GGATCC</u> CTA CAC CAC CTT CAC TTT GGT ACC) with KOD Hot Start DNA polymerase (Merck Millipore) according to manufacturer’s instructions. The resulting product was then cloned into the NdeI-BamHI site of the expression vector pET15b (Novagen) with the gene in-frame with the N-terminal His-tag to generate vector pET15b:SsuT3-His_tag. The TRD coding region of *modS* was then removed and the conserved 5′ and 3′ coding regions were fused. This was generated by using the pET15b::SsuT3-His_tag vector as a template via inverse PCR with primers SsuT3-TRD-remove-F (5′-CCCA AAT ATC TCA TCA CAG ATA GCT TTG AGG TTG G) and SsuT3-TRD-remove-R (5′-GCT GTT ATG CAG CTG AAT GCA GAA GAT GG) binding either side of the TRD, using KOD Hot Start DNA polymerase (Merck Millipore) according to manufacturer’s instructions. His-tagged TRD-less ModS was over-expressed in *E. coli* BL21 using isopropyl β-d-1-thiogalactopyranoside (IPTG) induction (0.5 mM) overnight at 37°C with 200 rpm shaking, and protein purified using TALON resin (Takara Bio) using standard protocols in 50 mM phosphate buffer pH7.4 containing 300mM NaCl.

### Generation of ModS Antisera

A cohort of five 6-8 week old female BALB/c mice (Animal Resources Centre, WA, Australia) were immunized subcutaneously with 25 μg of recombinant TRD-less ModS protein in 25uL PBS, mixed with 25 μL Freund’s Adjuvant (Merck, Darmstadt, Germany; Freund’s Complete Adjuvant (FCA) on day 0 and Freund’s Incomplete Adjuvant (FIA) subsequently on days 14, 21, 28 and 42. Terminal bleeds were collected on day 58 and serum separated via centrifugation. Pre-immune (naïve) serum was collected from cohorts prior to immunization via tail bleed. Serum was stored in 50% glycerol at −20°C. This antisera recognises all ModS alleles with equal affinity as it was raised against the conserved regions of ModS shared by all alleles. All animal work was carried out according to the Australian Code for the Care and Use of Animals for Scientific Purposes, with approval from the Griffith University Animal Ethics Committee (GLY/16/19/AEC).

### Single-Molecule, Real-Time (SMRT) sequencing and methylome analysis

Genomic DNA from our enriched triplet sets of *S. suis* strains LSS89 (*modS1* 19, 20, 21 repeats) and SS1056 (*modS2* 17, 18, 19 repeats) was prepared from an overnight culture in THB-Y broth and high-molecular-weight genomic DNA was isolated using the Sigma Genelute kit (Sigma Aldrich) according to the manufacturer′s instructions. SMRT sequencing and methylome analysis was carried out as previously described (46, 47). Briefly, DNA was sheared to an average length of approximately 5-10 kb (genomic DNA) using g-TUBEs (Covaris, Woburn, MA, USA) and SMRTbell template sequencing libraries were prepared using sheared DNA. DNA was end repaired, then ligated to hairpin adapters. Incompletely formed SMRTbell templates were degraded with a combination of Exonuclease III (New England Biolabs; Ipswich, MA, USA) and Exonuclease VII (USB; Cleveland, OH, USA). Primer was annealed and samples were sequenced on the PacBio Sequel system (Menlo Park, CA, USA) using standard protocols for long insert libraries. SMRT sequencing and methylome analysis was carried out at SNPSaurus (University of Oregon, USA).

### SWATH-MS proteomics

Overnight cultures of each *S. suis* strain (10^7^ CFU/ml) were harvested, lysed in guanidium buffer (6 M guanidium chloride, 50 mM Tris-HCl pH8, 10 mM dithiothreitol) and incubated at 30°C for 30 minutes with shaking (500 rpm). Cysteines of the total protein were alkylated by addition of acrylamide to a final concentration of 25 mM and incubated at 30°C for 60 minutes with shaking (500 rpm). Concentration of samples was assessed using a Nanodrop 2000 (Thermo Fisher). A 100 μg aliquot of the protein was then precipitated by addition of 1:1 methanol: acetone at −20°C overnight. The protein was pelleted at 18,000xg for 10 minutes and supernatant was removed before the pellet was resuspended in 50 μL trypsin reaction buffer and 1 μg trypsin (New England Biolabs) added and the suspension incubated overnight at 37°C. Tryptic digested peptides were then desalted and purified using a Ziptip (Millipore) as per manufacturer instructions. SWATH-MS was performed as previously described (48). Briefly, tryptic peptides were analyzed by LC-ESI-MS/MS using a Prominence nanoLC system (Shimadzu) and Triple TOF 5600 mass spectrometer with a Nanospray III interface (SCIEX). Peptides were separated on a Vydac EVEREST reversed-phase C18 HPLC column at a flow rate of 1 μL/min. A gradient of 10−60% buffer B over 45 min, with buffer A (1% acetonitrile and 0.1% formic acid) and buffer B (80% acetonitrile and 0.1% formic acid) was used. An MS-TOF scan was performed from an m/z range of 350− 1800 for 0.5 s followed by information dependent acquisition of MS/MS of the top 20 peptides from m/z 40−1800 for 0.05 s per spectrum, with automated CE selection. Identical LC conditions were used for SWATH-MS. SWATH-MS of triplicate biological replicates was performed with the same MS-TOF scan, followed by high sensitivity information-independent acquisition with m/z isolation windows with 1 m/z window overlap each for 0.1 s across an m/z range of 400−1250. Collision energy was automatically assigned by Analyst software (AB SCIEX) based on m/z window ranges. Proteins were identified by searching against *S. suis* Lss89 and SS1056 genomes (NCBI Accession GCA_900059105.1 and GCA_900051945.1 respectively) and common contaminants with standard settings using ProteinPilot 5.0.1 (AB SCIEX). False discovery rate analysis was performed on all searches. ProteinPilot search results were used as ion libraries for SWATH analyses. The abundance of proteins was measured automatically using PeakView (AB SCIEX) with standard settings. Comparison of protein relative abundance was performed based on protein intensities or ion intensities using a linear mixed-effects model with the MSstats package in R. Proteins with greater than X changes in abundance and with adjusted P-values. The mass spectrometry proteomics data have been deposited to the ProteomeXchange Consortium via the PRIDE (49) partner repository with the dataset identifier PXD023726.

### Minimum Inhibitory Concentration (MIC) assay

The MIC was measured by broth microdilution in triplicate experiments based on CLSI guidelines as described previously (50). Briefly, an overnight culture of *S. suis* was diluted to OD=0.1 (A_600_) and sub-cultured grown to mid log for 3 hours at 37°C. Mid log cultures were diluted to OD=0.2 (A_600_) and 50 μl of each culture was added to 96-well plates containing serially diluted antibiotic concentrations (5 μg/mL - 0.08μg/mL), and plates grown at 37°C with 5% CO_2_ for 24 hours. The MIC (mg/L) was determined as the last dilution at which turbidity was observed following overnight growth with all assays being performed in triplicate.

## Acknowledgements

We thank Amanda Nouwens and Peter Josh in SCMB MS facility, University of Queensland for conducting SWATH-MS proteomic analysis. We thank Allison Banse and Eric Johnson at SNPsaurus (University of Oregon, USA) for conducting SMRT sequencing and methylome analysis. This work was funded by an Australian Research Council Discovery Project DP180100976 to JMA and PJB, Australian National Health and Medical Research Council (NHMRC) Principal Research Fellowship 1138466 to MPJ.

## Supplementary figure legends

**Supplementary Figure 1. Expression of ModS is dependent on SSR tract length.** (A) Coomassie staining of whole cell lysates from enriched strains of S. suis. (B) Western blot using anti-ModS antisera demonstrates that ModS1 is only produced in strains enriched for 21 GAGCA repeats and that ModS2 is only produced in strains enriched for 18 repeats. Dotted line represents region of blot presented in Figure 1D.

**Supplementary Figure 2.** Three open reading frames occur in the *modS* gene due to variation in length of the GAGCA_(n)_ simple sequence repeat tract. Due to a frameshift down-stream of the GAGCA_(n)_ SSR tract, 19 and 20 repeats result in a premature stop codon (in bold, highlighted with *), and consequently no expression of the ModS protein; 21 GAGCA_(n)_ repeats in the SSR tract results in the gene being in-frame, and therefore ModS protein is produced.

## References

1. Goyette-Desjardins G, Auger JP, Xu J, Segura M, Gottschalk M. 2014. *Streptococcus suis*, an important pig pathogen and emerging zoonotic agent-an update on the worldwide distribution based on serotyping and sequence typing. Emerg Microbes Infect 3:e45.

2. Breton J, Mitchell WR, Rosendal S. 1986. Streptococcus suis in slaughter pigs and abattoir workers. Can J Vet Res 50:338–41.

3. Susilawathi NM, Tarini NMA, Fatmawati NND, Mayura PIB, Suryapraba AAA, Subrata M, Sudewi AAR, Mahardika GN. 2019. *Streptococcus suis*-Associated Meningitis, Bali, Indonesia, 2014-2017. Emerg Infect Dis 25:2235–2242.

4. Dutkiewicz J, Zajac V, Sroka J, Wasinski B, Cisak E, Sawczyn A, Kloc A, Wojcik-Fatla A. 2018. Streptococcus suis: a re-emerging pathogen associated with occupational exposure to pigs or pork products. Part II - Pathogenesis. Ann Agric Environ Med 25:186–203.

5. Tzeng YL, Thomas J, Stephens DS. 2016. Regulation of capsule in *Neisseria meningitidis*. Crit Rev Microbiol 42:759–72.

6. Atack JM, Winter LE, Jurcisek JA, Bakaletz LO, Barenkamp SJ, Jennings MP. 2015. Selection and Counterselection of Hia Expression Reveals a Key Role for Phase-Variable Expression of Hia in Infection Caused by Nontypeable Haemophilus influenzae. J Infect Dis 212:645–53.

7. Elango D, Schulz BL. 2020. Phase-Variable Glycosylation in Nontypeable *Haemophilus influenzae*. J Proteome Res 19:464–476.

8. Phillips ZN, Brizuela C, Jennison AV, Staples M, Grimwood K, Seib KL, Jennings MP, Atack JM. 2019. Analysis of Invasive Nontypeable *Haemophilus influenzae* Isolates Reveals Selection for the Expression State of Particular Phase-Variable Lipooligosaccharide Biosynthetic Genes. Infect Immun 87.

9. Seib KL, Srikhanta YN, Atack JM, Jennings MP. 2020. Epigenetic regulation of virulence and immunoevasion by phase-variable restriction-modification systems in bacterial pathogens. Annu Rev Microbiol 74:655–671.

10. Phillips ZN, Husna AU, Jennings MP, Seib KL, Atack JM. 2019. Phasevarions of bacterial pathogens - phase-variable epigenetic regulators evolving from restriction-modification systems. Microbiology 165:917–928.

11. Srikhanta YN, Maguire TL, Stacey KJ, Grimmond SM, Jennings MP. 2005. The phasevarion: a genetic system controlling coordinated, random switching of expression of multiple genes. Proc Natl Acad Sci U S A 102:5547–51.

12. Atack JM, Srikhanta YN, Fox KL, Jurcisek JA, Brockman KL, Clark TA, Boitano M, Power PM, Jen FE, McEwan AG, Grimmond SM, Smith AL, Barenkamp SJ, Korlach J, Bakaletz LO, Jennings MP. 2015. A biphasic epigenetic switch controls immunoevasion, virulence and niche adaptation in non-typeable Haemophilus influenzae. Nat Commun 6:7828.

13. Blakeway LV, Power PM, Jen FE, Worboys SR, Boitano M, Clark TA, Korlach J, Bakaletz LO, Jennings MP, Peak IR, Seib KL. 2014. ModM DNA methyltransferase methylome analysis reveals a potential role for Moraxella catarrhalis phasevarions in otitis media. FASEB J 28:5197–207.

14. Seib KL, Jen FE, Scott AL, Tan A, Jennings MP. 2017. Phase variation of DNA methyltransferases and the regulation of virulence and immune evasion in the pathogenic *Neisseria*. Pathog Dis 75.

15. Srikhanta YN, Gorrell RJ, Steen JA, Gawthorne JA, Kwok T, Grimmond SM, Robins-Browne RM, Jennings MP. 2011. Phasevarion mediated epigenetic gene regulation in *Helicobacter pylori*. PLoS One 6:e27569.

16. Srikhanta YN, Fung KY, Pollock GL, Bennett-Wood V, Howden BP, Hartland EL. 2017. Phasevarion-Regulated Virulence in the Emerging Pediatric Pathogen *Kingella kingae*. Infect Immun 85.

17. Brockman KL, Branstool MT, Atack JM, Robledo-Avila F, Partida-Sanchez S, Jennings MP, Bakaletz LO. 2017. The ModA2 Phasevarion of nontypeable Haemophilus influenzae Regulates Resistance to Oxidative Stress and Killing by Human Neutrophils. Sci Rep 7:3161.

18. VanWagoner TM, Atack JM, Nelson KL, Smith HK, Fox KL, Jennings MP, Stull TL, Smith AL. 2016. The modA10 phasevarion of nontypeable *Haemophilus influenzae* R2866 regulates multiple virulence-associated traits. Microb Pathog 92:60–67.

19. Gauntlett JC, Nilsson HO, Fulurija A, Marshall BJ, Benghezal M. 2014. Phase-variable restriction/modification systems are required for *Helicobacter pylori* colonization. Gut Pathog 6:35.

20. Phillips ZN, Tram G, Seib KL, Atack JM. 2019. Phase-variable bacterial loci: how bacteria gamble to maximise fitness in changing environments. Biochem Soc Trans 47:1131–1141.

21. Srikhanta YN, Dowideit SJ, Edwards JL, Falsetta ML, Wu HJ, Harrison OB, Fox KL, Seib KL, Maguire TL, Wang AH, Maiden MC, Grimmond SM, Apicella MA, Jennings MP. 2009. Phasevarions mediate random switching of gene expression in pathogenic *Neisseria*. PLoS Pathog 5:e1000400.

22. Atack JM, Yang Y, Seib KL, Zhou Y, Jennings MP. 2018. A survey of Type III restriction-modification systems reveals numerous, novel epigenetic regulators controlling phase-variable regulons; phasevarions. Nucleic Acids Res 46:3532–3542.

23. Atack JM, Weinert LA, Tucker AW, Husna AU, Wileman TM, F Hadjirin N, Hoa NT, Parkhill J, Maskell DJ, Blackall PJ, Jennings MP. 2018. *Streptococcus suis* contains multiple phase-variable methyltransferases that show a discrete lineage distribution. Nucleic Acids Res 46:11466–11476.

24. Katayama T, Kasho K, Kawakami H. 2017. The DnaA Cycle in *Escherichia coli:* Activation, Function and Inactivation of the Initiator Protein. Front Microbiol 8:2496.

25. Carabetta VJ, Tanner AW, Greco TM, Defrancesco M, Cristea IM, Dubnau D. 2013. A complex of YlbF, YmcA and YaaT regulates sporulation, competence and biofilm formation by accelerating the phosphorylation of Spo0A. Mol Microbiol 88:283–300.

26. Boddu RS, Perumal O, K D. 2020. Microbial Nitroreductases: A versatile tool for biomedical and environmental applications. Biotechnol Appl Biochem doi:10.1002/bab.2073.

27. Romero-Rodriguez A, Ruiz-Villafan B, Rocha-Mendoza D, Manzo-Ruiz M, Sanchez S. 2015. Biochemistry and regulatory functions of bacterial glucose kinases. Arch Biochem Biophys 577–578:1–10.

28. Schowanek D, Verstraete W. 1990. Phosphonate utilization by bacterial cultures and enrichments from environmental samples. Appl Environ Microbiol 56:895–903.

29. Schulze-Gahmen U, Pelaschier J, Yokota H, Kim R, Kim SH. 2003. Crystal structure of a hypothetical protein, TM841 of *Thermotoga maritima*, reveals its function as a fatty acid-binding protein. Proteins 50:526–30.

30. Asano Y, Lübbehüsen TL. 2000. Enzymes acting on peptides containing d-amino acid. Journal of Bioscience and Bioengineering 89:295–306.

31. Wei D, Wang M, Jiang B, Shi J, Hao J. 2014. Role of dihydroxyacetone kinases I and II in the dha regulon of *Klebsiella pneumoniae*. J Biotechnol 177:13–9.

32. Torrents E. 2014. Ribonucleotide reductases: essential enzymes for bacterial life. Front Cell Infect Microbiol 4:52.

33. Chan DI, Vogel HJ. 2010. Current understanding of fatty acid biosynthesis and the acyl carrier protein. Biochem J 430:1–19.

34. Gawthorne JA, Beatson SA, Srikhanta YN, Fox KL, Jennings MP. 2012. Origin of the diversity in DNA recognition domains in phasevarion associated modA genes of pathogenic *Neisseria* and *Haemophilus influenzae*. PLoS One 7:e32337.

35. dos Santos DF, Istvan P, Noronha EF, Quirino BF, Kruger RH. 2015. New dioxygenase from metagenomic library from Brazilian soil: insights into antibiotic resistance and bioremediation. Biotechnol Lett 37:1809–17.

36. Yongkiettrakul S, Maneerat K, Arechanajan B, Malila Y, Srimanote P, Gottschalk M, Visessanguan W. 2019. Antimicrobial susceptibility of *Streptococcus suis* isolated from diseased pigs, asymptomatic pigs, and human patients in Thailand. BMC Vet Res 15:5.

37. Lugsomya K, Chatsuwan T, Niyomtham W, Tummaruk P, Hampson DJ, Prapasarakul N. 2018. Routine Prophylactic Antimicrobial Use Is Associated with Increased Phenotypic and Genotypic Resistance in Commensal *Escherichia coli* Isolates Recovered from Healthy Fattening Pigs on Farms in Thailand. Microb Drug Resist 24:213–223.

38. Kouadio IK, Guessennd N, Dadie A, Koffi E, Dosso M. 2018. Comparative study of the impact of the administration of Amoxicillin and Algo-Bio((R)) alternative substance to antibiotics, on the level of selection of resistant *Enterobacteriaceae* in the digestive flora of piglets. J Glob Antimicrob Resist 13:161–164.

39. Callens B, Persoons D, Maes D, Laanen M, Postma M, Boyen F, Haesebrouck F, Butaye P, Catry B, Dewulf J. 2012. Prophylactic and metaphylactic antimicrobial use in Belgian fattening pig herds. Prev Vet Med 106:53–62.

40. Lekagul A, Tangcharoensathien V, Yeung S. 2019. Patterns of antibiotic use in global pig production: A systematic review. Vet Anim Sci 7:100058.

41. Krasny L, Bland P, Kogata N, Wai P, Howard BA, Natrajan RC, Huang PH. 2018. SWATH mass spectrometry as a tool for quantitative profiling of the matrisome. J Proteomics 189:11–22.

42. Brockman KL, Azzari PN, Branstool MT, Atack JM, Schulz BL, Jen FE, Jennings MP, Bakaletz LO. 2018. Epigenetic Regulation Alters Biofilm Architecture and Composition in Multiple Clinical Isolates of Nontypeable Haemophilus influenzae. mBio 9.

43. Atack JM, Guo C, Litfin T, Yang L, Blackall PJ, Zhou Y, Jennings MP. 2020. Systematic Analysis of REBASE Identifies Numerous Type I Restriction-Modification Systems with Duplicated, Distinct hsdS Specificity Genes That Can Switch System Specificity by Recombination. mSystems 5:e00497–20.

44. Atack JM, Guo C, Yang L, Zhou Y, Jennings MP. 2020. DNA sequence repeats identify numerous Type I restriction-modification systems that are potential epigenetic regulators controlling phase-variable regulons; phasevarions. FASEB J 34:1038–1051.

45. Weinert LA, Chaudhuri RR, Wang J, Peters SE, Corander J, Jombart T, Baig A, Howell KJ, Vehkala M, Valimaki N, Harris D, Chieu TT, Van Vinh Chau N, Campbell J, Schultsz C, Parkhill J, Bentley SD, Langford PR, Rycroft AN, Wren BW, Farrar J, Baker S, Hoa NT, Holden MT, Tucker AW, Maskell DJ, Consortium BRT. 2015. Genomic signatures of human and animal disease in the zoonotic pathogen *Streptococcus suis*. Nat Commun 6:6740.

46. Clark TA, Murray IA, Morgan RD, Kislyuk AO, Spittle KE, Boitano M, Fomenkov A, Roberts RJ, Korlach J. 2012. Characterization of DNA methyltransferase specificities using single-molecule, real-time DNA sequencing. Nucleic Acids Res 40:e29.

47. Murray IA, Clark TA, Morgan RD, Boitano M, Anton BP, Luong K, Fomenkov A, Turner SW, Korlach J, Roberts RJ. 2012. The methylomes of six bacteria. Nucleic Acids Res 40:11450–11462.

48. Peak IR, Chen A, Jen FE, Jennings C, Schulz BL, Saunders NJ, Khan A, Seifert HS, Jennings MP. 2016. *Neisseria meningitidis* Lacking the Major Porins PorA and PorB Is Viable and Modulates Apoptosis and the Oxidative Burst of Neutrophils. J Proteome Res 15:2356–65.

49. Perez-Riverol Y, Csordas A, Bai J, Bernal-Llinares M, Hewapathirana S, Kundu DJ, Inuganti A, Griss J, Mayer G, Eisenacher M, Perez E, Uszkoreit J, Pfeuffer J, Sachsenberg T, Yilmaz S, Tiwary S, Cox J, Audain E, Walzer M, Jarnuczak AF, Ternent T, Brazma A, Vizcaino JA. 2019. The PRIDE database and related tools and resources in 2019: improving support for quantification data. Nucleic Acids Res 47:D442–D450.

50. Peak IR, Jennings CD, Jen FE, Jennings MP. 2014. Role of *Neisseria meningitidis* PorA and PorB expression in antimicrobial susceptibility. Antimicrob Agents Chemother 58:614–616.

